# Prediction of best features in heterogeneous Lung adenocarcinoma samples using Least Absolute Shrinking and Selection Operator

**DOI:** 10.1101/765792

**Authors:** Ateeq Muhammed Khaliq, RG Sharathchandra, Meenakshi Rajamohan

## Abstract

This study aims to create a tumor heterogeneity-based model for predicting the best features of lung adenocarcinoma (LUAD) in multiple cancer subtypes using the Least Absolute Shrinking and Selection Operator (LASSO). The RNA-Seq raw count data of 533 LUAD samples and 59 normal samples were downloaded from the TCGA data portal. Based on consensus clustering method samples was divided into two subtypes, and clusters were validated using silhouette width. Furthermore, we estimated subtypes for the abundance of immune and non-immune stromal cell populations which infiltrated cancer tissue. We established the LASSO model for predicting each subtype’s best features. Enrichment pathway analysis was then carried out. Finally, the validity of the LASSO model for identifying features was established by the survival analysis. Our study suggests that the unsupervised clustering and Machine learning methods such as LASSO model-based feature selection can be effectively used to predict relevant genes which might play an essential role in cancer diagnosis.

## 1. Introduction

Lung cancer is reported to be the most deadly cancer [1]. Its shows the worst survival rate when compared with colon, breast, and pancreatic cancers combined. According to the American cancer society estimates both small cell and non-small cell lung cancer is the second most common cancer in both men and women. About 13% of all cancers are lung cancers. Lung Squamous cell carcinoma (LUSC) and Adenocarcinoma (LUAD) account for 15% and 85% of all lung cancer, respectively [2]. Lung cancer is a highly heterogeneous disease, and the identification of cancer subtypes is decisive for clinicians. Genetic mutations, cancer microenvironment, immune, and therapeutic selection pressures all dynamically contribute to tumor heterogeneity. Heterogeneity may lead to cells with a differential molecular signature within single tumor tissue, and in some cases, it may contribute to therapy resistance [3]. Therefore, deciphering LUAD heterogeneity will have a significant impact in designing precision medicine strategy. Heterogeneous data suffers from a large number of covariates, and identification of variable selection is necessary to obtain more accurate predictions with a large number of covariates.

Many computer-based diagnostic and predictive models have been used for predicting the risk of a variety of cancers, such as logistic regression, Cox proportional hazard model, artificial neural networks, decision trees and support vector machines. Previous studies indicate standard stepwise selection approaches which are not best for regression models with a vast number of covariates [4]. Alternatively, least absolute shrinkage and selection operator (LASSO), has received much attention for identification and selection of best variables. LASSO was introduced by Robert Tibshirani in 1996 [5]. Regularisation and feature selection are the two critical tasks LASSO performs. LASSO estimates the regression coefficients by maximising the log-likelihood function with the constraint that the sum of the absolute values of the regression coefficients, ∑j=1kβj, is less than or equal to a positive constant.

In this study, we selected the best features using LASSO established model. We downloaded the RNASeq data for LUAD samples from The Cancer Genome Atlas (TCGA) database. We differentiated the samples based on clusters into two subtypes to study the tumor heterogeneity. Differentially expressed genes (DEGs) were identified between two subtypes and normal groups, followed by predicting relevant variables that are associated with the response variable using the LASSO model and validating the variables using survival analysis. We also estimated the population abundance of tissue-infiltrating immune and stromal cell populations in each subtype to decipher the inflammatory, antigenic, and desmoplastic reactions occurring in cancer tissue. Our study provides new insight into tumor heterogeneity and its importance in sample classification for predicting biomarkers of LUAD.

## 2. Materials and Methods

### 2.1. Data source

The RNASeq data of LUAD, including 533 LUAD samples, and 59 normal samples were downloaded from the TCGA database (https://portal.gdc.cancer.gov/) in June 2019. All the raw, preprocessed data, images, supplemental tables and supporting files can be accessed at https://github.com/AteeqKhaliq/LUAD/.

### 2.2. Data preprocessing and grouping

533 Primary solid Tumor samples and 59 Solid Tissue Normal samples were downloaded from the TCGA database. We calculated a variance stabilising transformation (VST) from the raw count data and transformed the counts yielding a matrix of values approximately homoskedastic.

### 2.3. Molecular subtyping analysis

Feature dimension reduction was needed to remove irrelevant features and to reduce noises, and we used median absolute deviation (MAD) method and the features with MAD>0.5 were selected from set 2 groups. Consensus clustering (CC) [6] was used for the identification of subtypes on tumor samples. Silhouette width [7] was used to validate sample clustering.

### 2.4. Differential gene expression analysis

Differential gene expression (DGE) was assessed by using the DESeq2 package [8] (Version 1.24.0, https://bioconductor.org/packages/release/bioc/html/DESeq2.html) on Subtype-1 and Subtype-2 samples when compared with normal samples. Log2 fold change </> +/− 2 and P-value <0.05 were used as the cut-off values to identify the DEGs.

### 2.5. Construction of the LASSO model

Glmnet Package [9](Version 2.0-18, https://cran.r-project.org/web/packages/glmnet/index.html) was used to fit a generalised linear model via penalised maximum likelihood, LASSO model was established (Least Absolute Shrinkage and Selection Operator) on the DEGs from individual Subtype-1 and Subtype-2 cancer samples. We built a single pass (single fold) lasso-penalised model and performed 10-fold cross-validation to identify the best predictor.

### 2.6. Survival Analysis

To find clinically or biologically meaningful biomarkers Kaplan-Meier survival curves [10] were generated by selecting the best predictors from individual subtypes. Kaplan-Meier curves were generated using the TRGAted [11] (https://github.com/ncborcherding/TRGAted) package implemented in R.

### 2.7. Quantification of the abundance of the immune and stromal cell population in Cancer Subtypes

We estimated the abundance of tissue-infiltrating immune and non-immune stromal cell populations in Subtype-1 and Subtype-2 samples. MCP-counter [12] (https://github.com/ebecht/MCPcounter) Package was used to estimate the Microenvironment Cell Populations. VST normalised gene expression matrix was used for the estimation of an immune and stromal cell population in tumor samples.

### 2.8. Gene classification and enrichment analyses

clusterProfiler [13](Version 3.12.0, http://bioconductor.org/packages/release/bioc/html/clusterProfiler.html) was used to annotate the DEGs from Subtype-1 and Subtype-2 groups to biological processes, molecular functions, and cellular components in a directed acyclic graph structure with a q-value cutoff of 0.2, Kyoto Encyclopedia of Genes and Genomes (KEGG) [14] was utilized to annotate genes to pathways, and Disease Ontologies.

## 3. Results

### 3.1. Cancer subtype identification in LUAD samples

We used Consensus clustering (CC) method, an unsupervised clustering method for grouping subtypes in LUAD. CC method is the most widely used for subtype discovery in high dimensional datasets. We used settings of the agglomerative hierarchical clustering algorithm using Pearson correlation distance. Two distinct clusters were discovered in our datasets, 89 and 444 samples were clustered in Subtype-1 and Subtype-2 respectively (Fig.1.A). We have validated consistency within clusters of data using Silhouette width (Fig.1.B).

**Fig. 1.**
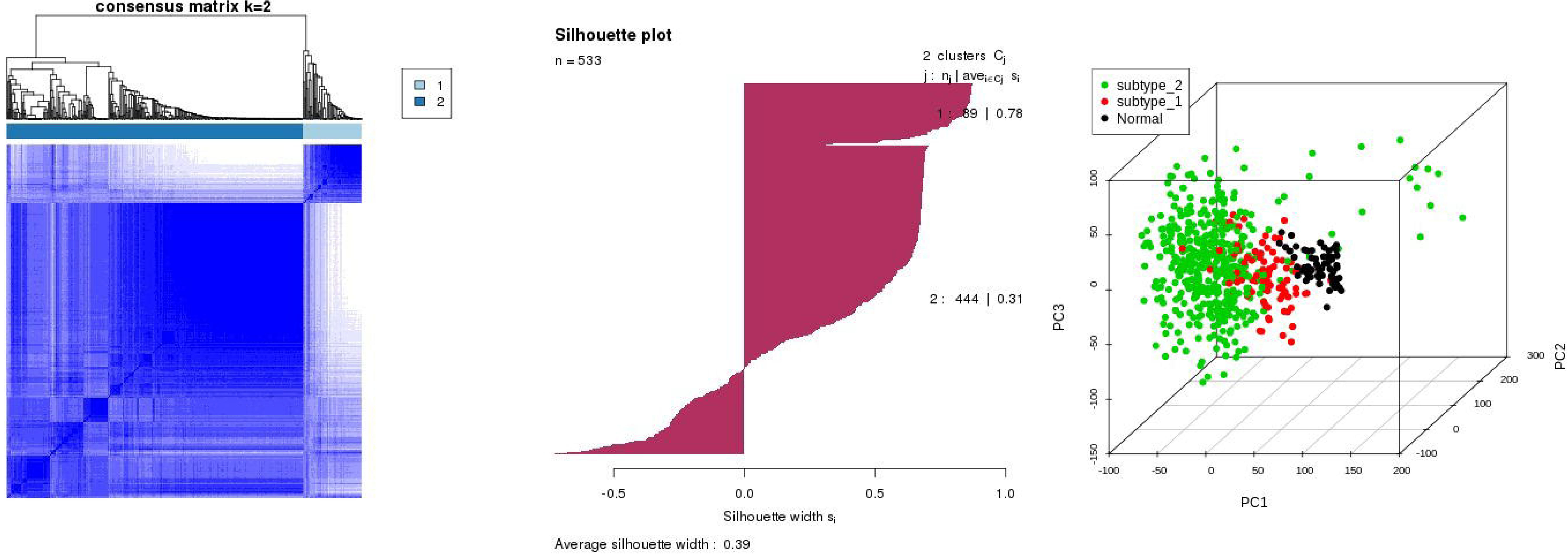
LUAD sample clustering. **(A)** CC plot shows the clustering of samples into two distinct subtypes. **(B)** Silhouette plot for validating the sample clustering. **(C)** PCA plot indicates distinct sample groups.

### 3.2. Identification of DEGs in Subtype-1 and Subtype-2 LUAD samples

We compared the subtype-1 and subtype-2 with the normal samples and based on the p-value cutoff < 0.05 and log2 fold change </> +/− 2 we identified significant DEGs. 2033 genes were upregulated, and 505 were downregulated in case of subtype-1 (Fig.2.A), and 5309 genes were upregulated, and 1219 were downregulated in case of subtype-2 (Fig.2.B) shows differential expression pattern in subtype-1 and subtype-2. The DEGs in both subtypes were used for building the LASSO predictive model and for the identification of best predictor genes in LUAD heterogeneous cancer data.

**Fig. 2.**
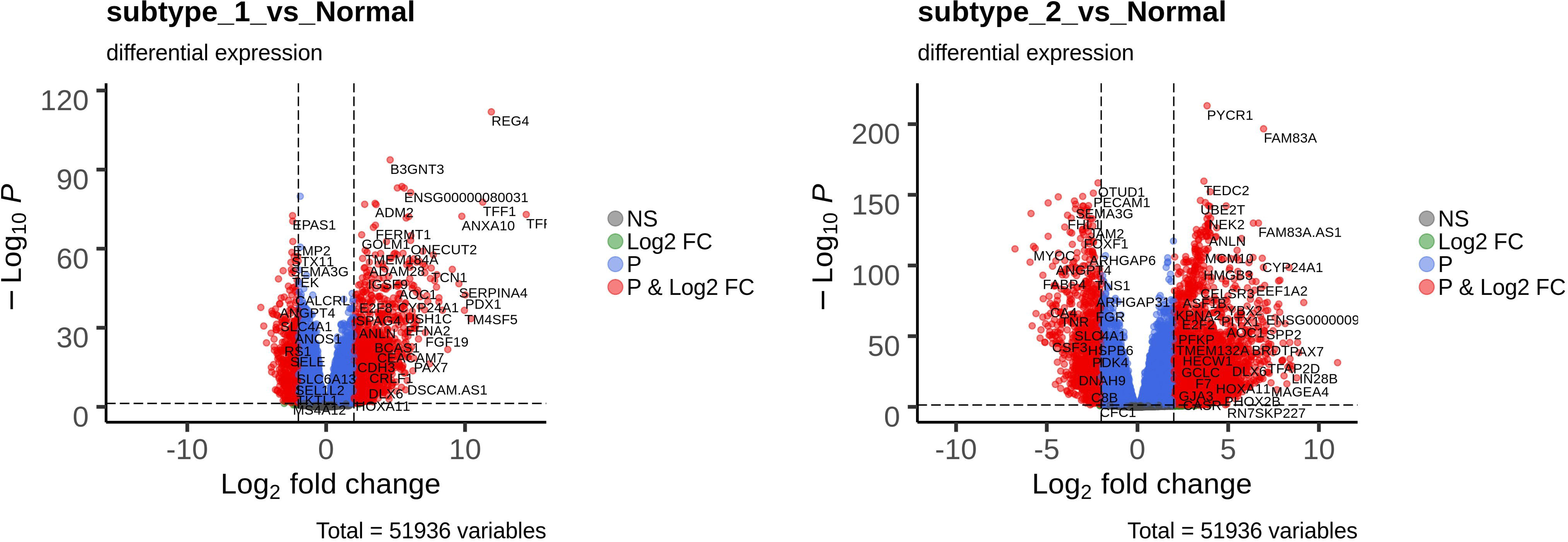
DGE analysis at standard cutoff of </>+/− 2 fold change at p value of <0.05. (A) DGE volcano plot for Subtype-1 samples. (B) DGE volcano plot for Subtype-2 samples.

### 3.3. LASSO model for identification of best predictive genes

LASSO was first described by Robert Tibshirani in 1996. Regularisation and feature selection are the two crucial task performed by LASSO. RNASeq datasets are high dimensional datasets, with smaller sample size and a large number of features also called small-n-large-p datasets (p ≫ n). High dimensional data will be sparse, and only a few features affect the response variable, and LASSO is known to identify the best features that affect the response variable. We deal with a p ≫ n situation for feature selection in our Subtype-1 and Subtype-2 datasets, thus probably not all DEGs are relevant for the identification of features which affect the response variable. The purpose of our analysis is to identify the feature selection task and underline which genes are more relevant to predict and to classify them as biomarkers, to do so, we have used the LASSO model.

The result shows the trends of the 40 and 43 most relevant features selected by our model in subtype-1 and subtype-2 LUAD, respectively. The next step would be to find the most appropriate values for λ for our LASSO model. We analysed the λ value using the 10 fold cross-validation, between λ min that gives minimum mean cross-validated error or λ1se, that gives a model such that error is within one standard error of the minimum. Using this analysis, we obtained the most relevant genes which are unique to subtype-1 and subtype-2 in the detection of a LUAD. A list of best-predicted genes available for each cancer subtypes is shown in Table 1.

### 3.4. Analysis of the microenvironment of Subtype-1 and Subtype-2 LUAD samples

The abundance of tissue-infiltrating immune and non-immune stromal cell populations is highly informative. It has been shown that the extent of infiltrating immune cells is associated with disease prognosis. T-cell infiltrates, endothelial cells and fibroblasts are associated with a favorable outcome and also poor prognosis in some cancer types [15]. To understand the immunological microenvironment in our expression subset-1 and subset-2 we used MCP-counter method as described by Becht et al.[12]. The estimations consist of single sample scores which are computed on each sample independently in two subtypes. The heatmap shown in Figure 3 clearly distinguish our subtype-1 and subtype-2 into two different categories based on tissue-infiltrating immune and non-immune stromal cell populations. Subtype-1 shows apparent increase in B lineage cells, monocytic cells, Cytotoxic lymphocytes, Natural killer cells, and CB 8 T cells and Subtype-2 shows decreased levels of T-cells, macrophages, B cells, and natural killer (NK) cells, as well as endothelial cells and fibroblasts. Our study clearly distinguishes LUAD subtypes based on their inflammatory and stromal profiles, and Subtype-1 LUAD samples show increased expression of immunological markers than Subtype 2 samples.

**Fig. 3.**
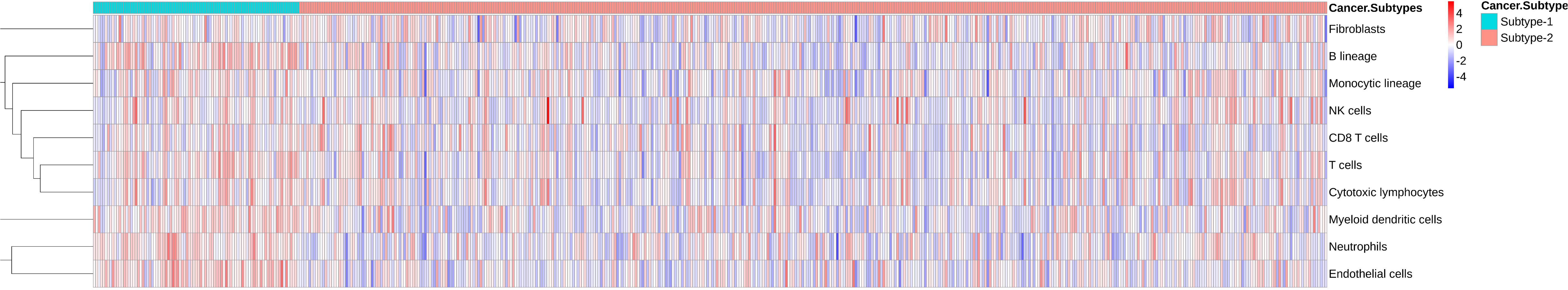
The heatmap distinguish subtype-1 and subtype-2 into two different categories based on tissue-infiltrating immune and non-immune stromal cell population.

### 3.5. Pathway analysis

Subtype-1 and Subtype-2 showed distinct and unique features which are involved in cancer progression. Genes such as IL22RA2, PLA2G2C identified in subtype1 are upregulated and found to be involved in canonical cancer regulatory pathways such as the JAK-STAT signalling pathway and RAS signalling pathway (Fig. 4). The PLA2G2C gene is associated with alpha-Linolenic acid as well as ether lipid metabolism, and both are known to play a role in cancer progression [16]. Our model identified ONECUT1 gene in subtype1, which is associated with regulating pluripotency of stem cells and DEFA3 gene which is associated with Transcriptional misregulation in cancer. The gene AWAT2 identified in subtype1 is found to be involved in Retinol metabolism, and studies suggest that retinoid signalling triggers tumor development [17]. The NECTIN4 gene highlighted in our study is associated with Adherens junction, which plays an essential role in cancer initiation and progression [18]. PYCR1 gene involved in proline metabolism which is highly correlated with cancer is also found to be differentially expressed in subtype1. EFNA3 is upregulated in Subtype-2, and previous studies show that it contributes to the metastatic spread of breast cancer [19]. Our model predicted that increased expression of LGR4, TESC, TOP2A, and ZNF695 is suggestive of increased invasive and metastatic activity in Subtype-2 [20]. Decreased expression of FGF10 is seen in Subtype-2 and predicted by our model is suggestive of dysplasia in LUAD subtype-2 samples [21].

**Fig. 4.**
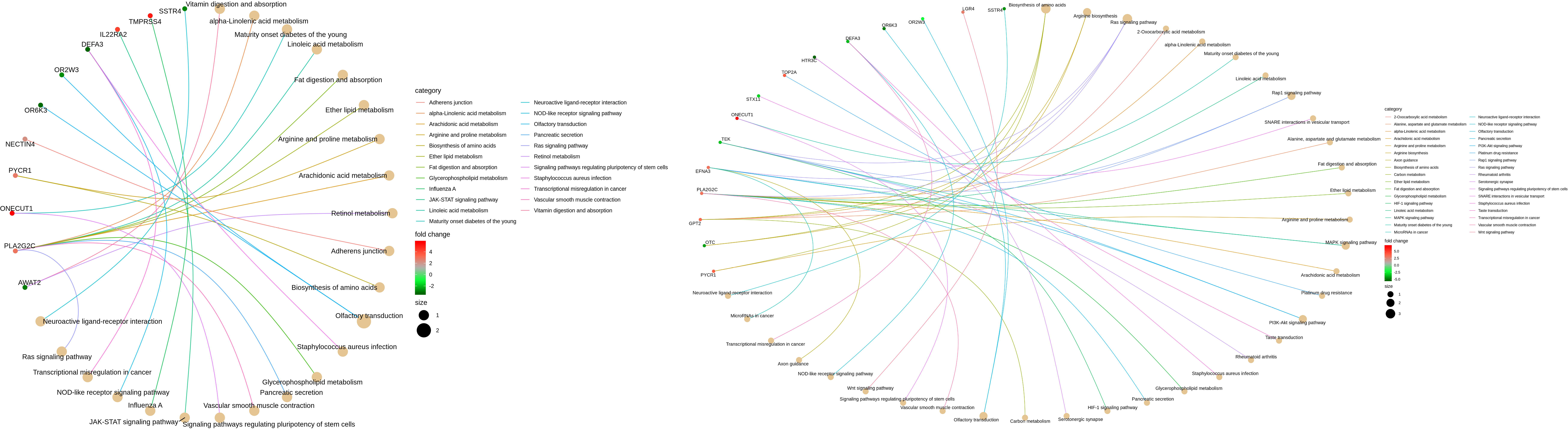
Pathway analysis for the best predicted genes by LASSO model. (A) Subtype-1 LASSO predicted genes pathway analysis. (B) Subtype-2 LASSO predicted genes pathway analysis.

### 3.6. Validation with survival analysis

The LUAD samples were classified into two subtypes based on the consensus clustering method. Overall survival analysis for the most predictive genes identified by our model in Subtype-1 and Subtype-2 groups was conducted. The genes predicted by our LASSO model accurately predicted the outcome of a patient’s survival using gene expression data. Genes such as LAD1, NECTIN4, SLC25A48, LYVE1, EFNA3, FGF10, HELT, HTR3C, and LGI3 yielded accurate predictions for the risk of LUAD and can be used in cancer prediction. Survival plots and its p-value is shown in Supplemental Figures.

## 4. Discussion

In this study, we developed a LASSO based model for accurate feature selection in LUAD. Our model removed variables that are redundant and removed features which do not add any valuable information in disease prediction. Analysis using the survival data for the predicted genes showed that the model could effectively predict genes responsible for disease prognosis in high dimensional datasets. Deciphering cancer heterogeneity is very critical in understanding cancer dynamics and also for the development of personalised cancer treatment [22]. We used Consensus clustering method to determine the number of clusters in our samples, and we clustered the samples into two groups which produced optimal silhouette width for the determined clusters. Differential gene expression analysis showed distinct expression patterns in both Subtype-1 and Subtype-2.

The number of differentially expressed genes was very high, and in these situations, it is difficult to predict the relevant variables. LASSO model was established around DE genes in Subtype-1 and Subtype-2 groups. Not all the expressed genes were relevant, and our model predicted the most relevant genes which were involved in disease progression. Decreased expression of LYVE1 and MED28P8 in Subtype-1 and FGF10, HELT, HTR3C, LGI3, PACRG, PLAC9P1, and STX11 in Subtype-2 showed worse overall survival in LUAD samples. Whereas increased expression of genes such as CDC37P2, DCST1, IL22RA2, LAD1, NECTIN4, SLC25A48, TMEM51.AS1, TMPRSS4, VPS9D1-AS1, AWAT2, BCL9P1 and CD5L in Subtype-1 and EFNA3, LGR4, TESC, TOP2A and ZNF695 in subtype-2 showed decreased overall survival in LUAD samples.

Long intervening noncoding RNAs (lncRNAs) are known to be critical regulators of numerous biological processes, and substantial evidence supports that lncRNA expression plays a significant role in tumorigenesis and tumor progression. Increased expression of LINC00862 in Subtype-1 samples correlates with worse survival in LUAD subtypes. Whereas, decreased expression of LINC00211 in Subtype-1 and LINC01506, LINC01785 and LINC01996 in Subtype-2 showed worse survival in LUAD samples. Our LASSO model predicts the most relevant and distinct genes from Subtype-1 and Subtype-2 samples which might be an important factor in cancer diagnosis and management.

The best predictors for subtype-1 and subtype-2 from the LASSO model were found to be involved in several regulatory pathways. Subtype-1 gene such as S100A12 is a vital serum inflammatory marker and has been illustrated in several cancer types such as oropharyngeal squamous cell carcinoma and gastric cancers [23]. Subtype-1 samples show increased expression of CD5L and TMPRSS4, which induces cancer stem cell-like properties and promotes malignant transformation by limiting lung epithelial cell apoptosis and promoting immune escape in NSCLC patients [24]. Long noncoding RNA VPS9D1-AS1 overexpression in subtype-1 predicts poor prognosis and serves as a biomarker to predict the prognosis of NSCLC [25]. Overexpression of nectin-4 oncoprotein, LYVE-1/PCAB, PROM2, and LAD1 are associated with poor overall survival in subtype-1 samples and can be considered as candidate serum and tissue biomarker as well as therapeutic target [26].

Overexpression of Lgr4 in Subtype-2 samples promotes tumor aggressiveness may potentially become a novel target for cancer therapy [27]. Subtype-2 samples show upregulation of TOP2A and ZNF695 is associated with worse prognosis and induces overrepresentation of growth and proliferation pathways and can act as prognostic and predictive markers [28]. Hypoxia-inducible oncogene EFNA3 is overexpressed in subtype-2 samples may play a critical role in the focal adhesion kinase (FAK) signalling and VEGF-associated tumor angiogenesis pathway [19]. Downregulation of tumor suppressor gene STX11 in Subtype-2 samples predicts poor prognosis. Various studies indicate the role of STX11 expression in suppressing the proliferation of T-cell [29].

Our model predicts lncRNAs such as LINC01506, LINC01785, LINC01996, LINC00862, and LINC02014 were expressed in subtype-1 and subtype-2 samples which can be considered as potential biomarkers and shows poor overall survival in LUAD. LncRNAs might be used as biomarkers and drug targets for early diagnosis, prognosis and personalised treatment of LUAD patients.

Our study suggests that Consensus Clustering methods and LASSO combined will help us to develop a model with the most appropriate characteristics. Consistent with these finding, different subtypes showed distinct unique features which underscore the importance of sample grouping and assessment. Furthermore, Survival analyses validate that the survival time of the predicted genes correlates with gene expression pattern, which is recognisably different in both the Subtypes, indicating that LASSO model could effectively be used to overcome the feature selection problem and can be used for accurate prediction of risk in LUAD.

## Funding

This research did not receive any specific grant from funding agencies in the public, commercial, or not-for-profit sectors.

## Table description

**Table 1:** List of best-predicted genes by LASSO Model for cancer Subtype-1 and Subtype-2

## Notes

https://github.com/AteeqKhaliq/LUAD/

